# A Novel Role for Phospholamban in the Thalamic Reticular Nucleus

**DOI:** 10.1101/2023.11.22.568306

**Authors:** Benjamin Klocke, Aikaterini Britzolaki, Joseph Saurine, Hayden Ott, Kylie Krone, Kiara Bahamonde, Connor Thelen, Christos Tzimas, Despina Sanoudou, Evangelia G. Kranias, Pothitos M. Pitychoutis

## Abstract

The thalamic reticular nucleus (TRN) is a critical brain region that greatly influences vital neurobehavioral processes, including executive functioning and the generation of sleep rhythms. Recently, TRN dysfunction was suggested to underlie hyperactivity, attention deficits, and sleep disturbances observed across various devastating neurodevelopmental disorders, including autism, schizophrenia and attention-deficit/hyperactivity disorder (ADHD). Notably, a highly specialized sarco- endoplasmic reticulum calcium (Ca^2+^) ATPase 2 (SERCA2)-dependent Ca^2+^ signaling network operates in the dendrites of TRN neurons to regulate their high-frequency bursting activity. Phospholamban (PLN) is a prominent regulator of the SERCA2 with an established role in maintaining Ca^2+^ homeostasis in the heart; although the interaction of PLN with SERCA2 has been largely regarded as cardiac-specific, our findings challenge this view and suggest that the role of PLN extends beyond the cardiovascular system to impact brain function. Specifically, we found PLN to be expressed in the TRN neurons of the adult mouse brain and utilized global constitutive and innovative conditional genetic mouse models, in combination with 5-choice serial reaction time task (5-CSRTT) and electroencephalography (EEG)-based somnography to assess the role of PLN in regulating executive functioning and sleep, two complex behaviors that map onto thalamic reticular circuits. Overall, the results of the present study show that perturbed PLN function in the TRN results in aberrant thalamic reticular behavioral phenotypes in mice (i.e., hyperactivity, impulsivity and sleep deficits) and support a novel role for PLN as a critical regulator of the SERCA2 in the thalamic reticular neurocircuitry.

## 1. BACKGROUND

The thalamic reticular nucleus (TRN) is a critical but poorly understood structure in the mammalian forebrain involved in regulating complex behaviors, including sleep and executive functioning [1]. The TRN is uniquely situated between the cortex and the thalamus, and consists of a thin layer of gamma-aminobutyric acid (GABA)-ergic neurons that inhibit thalamic relay cells [2]. Interestingly, the activity of TRN neurons relies heavily on calcium (Ca^2+^) regulation; depending on their resting membrane potential, TRN neurons fire in two distinct modes; at depolarized states, they fire tonic sodium (Na^+^) spikes, but when hyperpolarized they generate repetitive “low-threshold” Ca^2+^ transients crowned by high-frequency Na^+^ spikes known as bursts [3–5]. Notably, a highly specialized sarco-(SR)-endoplasmic reticulum (ER) Ca^2+^ ATPase2 (SERCA2)-dependent Ca^2+^ signaling network operates in the dendrites of TRN neurons to generate and regulate the strength of their high-frequency bursting activity [3, 6].

In mammals, three differentially expressed SERCA genes (known as *ATP2A1-3* in humans) encode at least ten SERCA isoforms [7]. Of all SERCA proteins, SERCA2 is the most widely distributed, with the SERCA2b isoform being ubiquitously expressed in nerve cells [8]. Given its critical role in ensuring appropriate ER Ca^2+^ store filling [9, 10], perturbed function of the SERCA2 may result in aberrations in intracellular Ca^2+^-signaling cascades that may in turn impact neuronal function [11]. Indeed, SERCA2-dependent dysregulation of neuronal Ca^2+^ homeostasis has been implicated in the pathophysiology of devastating disorders that affect cognition, including schizophrenia, Alzheimer’s disease, and Darier’s disease [12–17]. Of note, Phospholamban (PLN) is a critical regulator of the SERCA2 with an established role in maintaining Ca^2+^ homeostasis in cardiac, skeletal and smooth muscle cells [18–20]. Specifically, in myocardial cells dephosphorylated PLN binds to SERCA2 and reduces its affinity for Ca^2+^, thus decreasing Ca^2+^ uptake into the SR [18]. The role of PLN in SR Ca^2+^ cycling and cardiomyocyte contractility is well defined; indeed, increased PLN inhibition of SR Ca^2+^ cycling is considered a major characteristic of heart failure in humans [18, 21–24]. Notably, constitutive genetic ablation of PLN in mice (*i.e.*, *Pln^-/-^*) results in phenotypically normal animals that have served as an invaluable model system for elucidating the Ca^2+^-handling role of PLN in myocardial cells [25–27].

Even though SERCA2 is ubiquitously expressed in neurons [8, 28, 29], a number of early Northern blotting studies had failed to detect *Pln* mRNA expression in the brain of humans or mice [30, 31]. Excitingly, we found that the *Pln* gene is selectively expressed at the protein level in the mouse GABAergic TRN neurons, and thus hypothesized that PLN may be necessary for modulating critical behaviors that map onto thalamic reticular circuits. In order to address this hypothesis, we employed a combination of existing global (i.e., *Pln^-/-^*) and novel conditional genetic knockout mouse models. Given the TRN-specific expression profile of PLN, we developed a novel GABAergic Cre- conditional *Pln* knockout mouse line (cKO) in which PLN expression is selectively ablated in the GABAergic TRN neurons, and further assessed how PLN deficiency in the TRN impacts locomotor activity, executive functioning and sleep architecture, three vital behaviors that map onto thalamic reticular circuits. Overall, our findings provide the first evidence that the PLN/SERCA2-mediated Ca^2+^ signaling is critical for information gating by the TRN and is required for appropriate elaboration of sleep, cognitive and locomotor behaviors.

## 2. MATERIALS AND METHODS

### 2.1. Animals

Animals were bred in the animal facility of the University of Dayton (Dayton, OH) under standard 12h light/dark schedule and housed in groups of 3-4 of the same genetic background and sex after weaning, unless otherwise stated in standard transparent mouse cages (300mm x 160mm x110 mm; Alternative Design, AR, USA). Adult (*i.e*., 8-12 month old) male and female *Pln^-/-^* and wild type *Pln^+/+^* 129SvJ littermate mice derived from breeders provided by Dr. Evangelia G. Kranias (University of Cincinnati, Cincinnati, OH, USA; Biomedical Research Foundation of the Academy of Athens, Athens, Greece) [32, 33]. The generation of the new floxed PLN mouse line is described below. Mice were housed under standard conditions (i.e., 22°C ± 2°C; 30-60% humidity; 12-h light/dark cycle, lights on at 7:00 AM), and given access to food and water *ad libitum*, unless otherwise stated. Mouse genotypes were independently determined in littermates derived from heterozygote crosses using standard genotyping procedures. All animals were habituated to the experimental conditions for at least one week prior to any experiments, while behavioral testing occurred during the light cycle, unless otherwise stated. All experiments were conducted in accordance with the National Institute of Health (NIH) Guide for the Care and Use of Laboratory Animals (NIH Publications No. 80-23; revised 1978) and were approved by the University of Dayton Animal Care and Use Committee (IACUC).

### 2.2. Construction of the *Pln* LoxP-Cre conditional knockout mouse line (cKO)

The construction of the floxed *Pln (i.e, Pln^lox/lox^*) mouse line was funded by an inaugural STEM Catalyst grant from the University of Dayton to Dr. Pitychoutis and service was provided by the Transgenic Animal and Genome Editing Core at Cincinnati Children’s Hospital Medical Center (CCHMC; Cincinnati, OH) **(Fig. 1)**. Specifically, the floxed *Pln* allele was generated by CRISPR/Cas9 using a dual-sgRNA strategy to cut the genomic DNA fragment containing the entire coding sequence and repair it with a sgRNA-resistant long ssDNA donor with two loxP sites, the left homologous arm at 76nt and the right at 109nt, the so-called Easi-CRISPR method [34] **(Fig. 1a)**. The sgRNA target sequences were selected according to the on- and off-target scores from the web tool CRISPOR (http://crispor.tefor.net) [35]. sgRNAs were transcribed *in vitro* using the MEGAshorscript T7 kit (ThermoFisher) and purified by the MEGAclear Kit (ThermoFisher) and stored at -80°C. To validate their activity, we incubated sgRNAs and Cas9 protein (IDT) at 37°C for 10 min to form ribonucleoproteins (RNP). The final concentrations are 50 ng/ul of each sgRNA and 150 ng/ul Cas9 protein. The RNP was delivered into a small batch of mouse zygotes by electroporation using a Genome Editor electroporator (BEX; 30V, 1ms width, and 5 pulses with 1s interval). Zygotes were cultured for 4 days to blastocysts and genotyped individually, where we found that 50% embryos showed the large deletion between two cut sites. The long ssDNA donor was produced in-house by releasing it from a dsDNA plasmid by double nickase digestion (Nb.BsmI), followed by denaturing in the RNA-loading buffer at 70°C for 5 min then on ice for 1 min, electrophoresis, and gel extraction with QIAquick Gel Extraction Kit (Qiagen). The mosaic floxed mice were created by electroporating zygotes on the C57BL/6 genetic background with the RNP and long ssDNA donor at the concentration of 50 ng/uL. Electroporated zygotes were transferred into the oviductal ampulla of pseudopregnant CD-1 females on the same day. Pups were born and genotyped by PCR and Sanger sequencing. One mouse *(i.e.*, Mouse #5040) containing the LoxP sites was then selected as confirmed by primer pairs P1-P3 and P4-P6 for 5’ and 3’ end **(Fig. 1b,c)**. The selected floxed PLN mosaic female mouse was backcrossed to a wild type male mouse; the resulting pups were genotyped by PCR and Sanger sequencing to confirm the presence of both 5’- and 3’-LoxP sites **(Fig. 1d)**. Upon expansion of the colony, homozygote floxed PLN mice were crossed with another transgenic mouse line expressing the Cre recombinase in GABAergic GAD65^+^ neurons, *Gad2-Cre^+/+^;Pln^+/+^* (*i.e.,* B6J.Cg- *Gad2*^tm2(cre)Zjh^/MwarJ mice; The Jackson Laboratory, stock No. 028867). Floxed PLN heterozygotes carrying the Cre recombinase were selected with PCR genotyping and crossed again with *Pln^lox/lox^*mice to generate litters containing the *Gad2-Cre^+/0^;Pln^lox/lox^*(cKO) and their *Gad2-Cre^0/0^; Pln^lox/lox^* (Control) littermates **(Fig. 1e)**.

**Figure 1:**
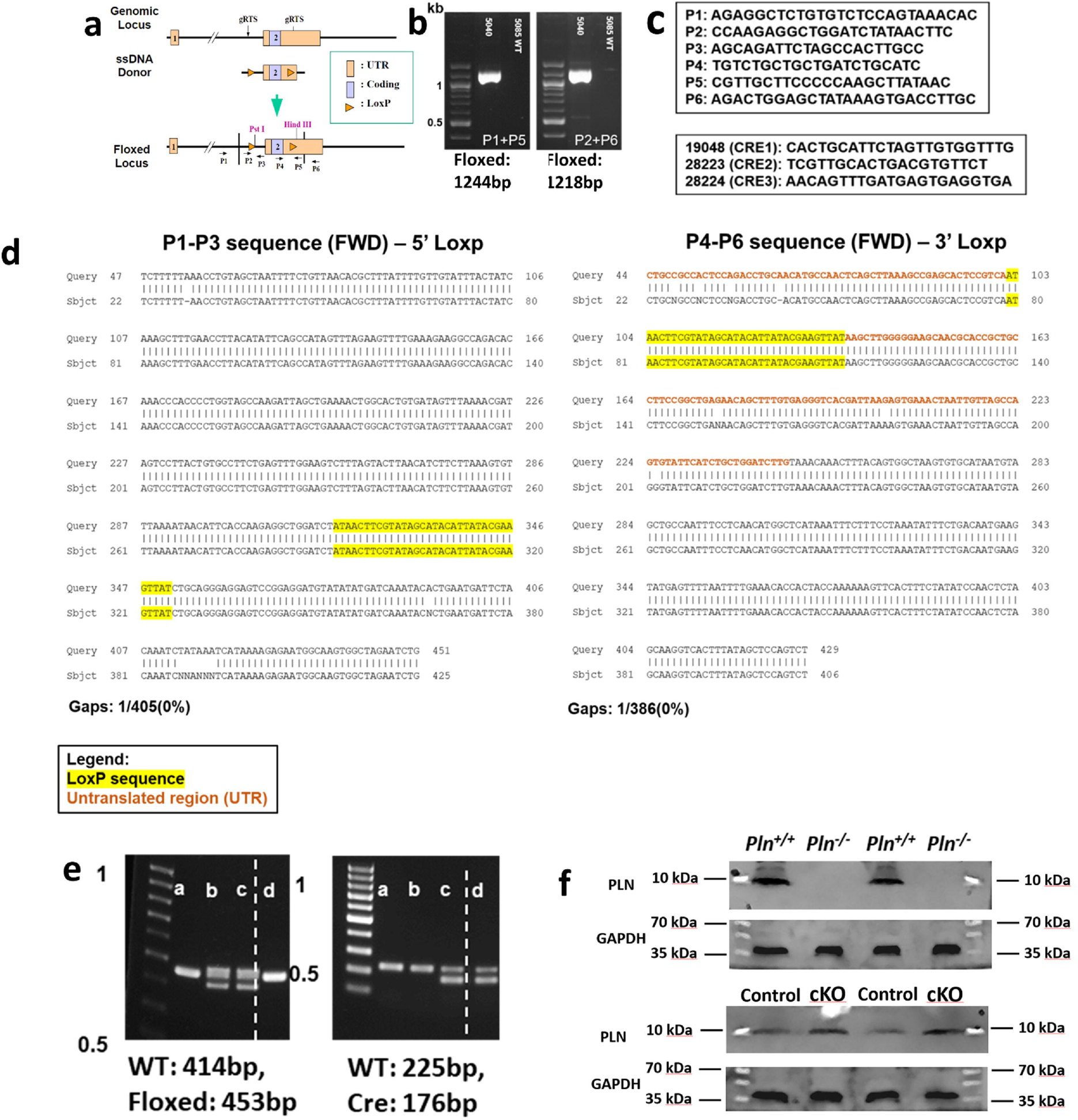
Construction of the *Pln* LoxP-Cre conditional knockout mouse line (cKO): **a)** schematic illustration of the targeting strategy: two LoxP sequences were inserted upstream and downstream of the *Pln* gene using a dual-sgRNA strategy to cut the genomic DNA fragment containing the entire coding sequence and repair it with a sgRNA-resistant long ssDNA (Easi-CRISPR method). CRISPR reagents were electroporated into zygotes and successful embryos were then transferred into the oviductal ampulla of pseudopregnant CD-1 females. The final floxed *Pln* locus contains two LoxP sequences flanking the *Pln* coding region and two restriction enzyme sites (PstI, HindIII); **b)** the mosaic floxed PLN mouse (#5040) was selected after confirming the presence of both LoxP sites; **c)** primers used for the genotyping of the *Pln^lox/lox^*, control and cKO mice. **d)** Representative Sanger sequencing results confirming the integration of both LoxP sites in *Pln^lox/lox^* mice; **e)** representative PCR genotyping blots of littermates derived from the final crossing that generated the cKO mice; cKO mice were floxed homozygotes (control) and heterozygote to Cre (*i.e., Gad2-Cre^0/+^*); control mice were floxed homozygotes (control) without carrying the Cre (*i.e., Gad2-Cre^0/0^*). **f)** Conditional deletion of PLN in GABAergic TRN neurons was targeted to the brain and did not result in diminished PLN protein expression in the heart of cKO mice, as assessed by western blotting.

### 2.3. PLN mRNA and Protein Expression Assessments

Total RNA was isolated from snap-frozen whole mouse brain with the Trizol protocol (Life Technologies) and cDNA was synthesized with the Superscript-RT kit (Invitrogen) using oligo-dT primers. PLN mRNA expression in the whole mouse brain was assessed using RT-PCR procedures and the following primers: Forward T1T2: CCCATAAACCTGGGAACAGA; Reverse T1T2: TAGCCGAGCGAGTGAGGTAT, yielding a transcript variant of 235bp. PLN protein expression was detected in the mouse heart using standard immunoblotting procedures. Briefly, whole heart tissue was collected from *Pln^+/+^, Pln^-/-^,* Control and cKO mice and sonicated in RIPA buffer (150nM NaCl, 1.0% Triton X-100, 0.5% sodium deoxycholate, 0.1% SDS, 50mM Tris, pH 8.0). Total protein quantification was assessed via the BCA assay (Pierce BCA Protein Assay Kit #23227). For each sample, equal amounts of protein (i.e., 40ug) was loaded onto a 10% SDS-PAGE gel for electrophoresis. Proteins were transferred onto a nitrocellulose membrane and probed with primary PLN (Invitrogen #MA3-922; 1:500) and GAPDH (Novus Biologicals #NB100-56875; 1:200) antibodies overnight (O/N) at 4०C. Upon incubation with alkaline phosphatase-conjugated secondary antibody (Invitrogen #A16014, #A16002; 1:5,000), membranes were exposed to ECF substrate (GE Healthcare #1067873) for visualization in a UVP BioSpectrum 500 Imaging System.

### 2.4. Free-floating fluorescent immunohistochemistry (IHC) for detection of PLN immunoreactivity

Anesthetized mice were transcardially perfused with ice-cold 1xPBS followed by 4% paraformaldehyde (PFA) fixative solution. Isolated mouse brains were then post-fixed in 4%PFA O/N at 4 °C prior to sectioning using a Vibratome (Lafayette Instruments; IN, USA). 40μm free-floating coronal Vibratome-cut sections were first incubated in a 10% normal goat serum (NGS) blocking solution in 1x TBS-0.1% Triton X-100 for 1 hour at room temperature (RT) and were then incubated O/N at 4°C in primary antibody solution in 3% NGS. The next day sections were washed 3x15min in 1xTBS, incubated in blocking solution for 1h at RT, and were then incubated for 1h at RT in the secondary antibody solution in 3% NGS in the dark. Following 3x15min in 1xTBS washes, sections were mounted onto positively charged glass slides and coversliped using DAKO fluorescent mounting medium (Agilent Technologies S3023). A negative control (i.e., sections incubated in 3% NGS solution in the absence of primary antibody solution) was included in each IHC experiment. Images were taken with an Olympus Fluoview 3000 confocal microscope (PA, USA). Brain sections from the different mouse genotypes were processed concurrently on each IHC experiment, while imaging conditions (i.e., laser power, HV, gain and offset) were kept consistent throughout the samples within the same brain region and magnification of the respective experiment. Image processing (*i.e.*, brightness and contrast), if needed, was kept to minimum and was equally applied to all images. The primary antibodies used in our single and/or double IHC experiments included mouse monoclonal PLN (Invitrogen #MA3-922; 1:100), rabbit polyclonal anti-GAD67 (Invitrogen #PA5-21397; 1:250), rabbit polyclonal anti-SERCA2 (Abcam #ab150435; 1:500), and rabbit polyclonal anti-GFAP (Abcam #ab7260, 1:500) and secondary antibodies goat anti-mouse Alexa Fluor 488 (Invitrogen #A32723TR; 1:500), goat anti mouse IgG2a Alexa Fluor 647 (Invitrogen #A21241, 1:500), and goat anti-rabbit Alexa Fluor 568 (Invitrogen #A21428; 1:500). IHC immunoreactivity for PLN, GAD-67 and SERCA2 was first assessed in single IHC experiments, and their co-localization in TRN neurons was further confirmed in double IHC experiments. The specificity of the primary PLN Ab used for IHC experiments was also validated using western blotting for the detection of PLN in the heart of *Pln^-/-^* and *Pln^+/+^* mice **(Fig. 1f).**

### 2.5. Behavioral testing

#### 2.5.1. Open field test (OFT)

Spontaneous locomotor activity and anxiety-like behavior were assessed by individually placing animals in the center of a novel OFT arena (45cm x 45cm x 40cm). Both the horizontal distance traveled, and the time mice spent in the center of the arena were recorded and analyzed using the SMART v3.0 video tracking software (Panlab, Coulbourn Instruments, Harvard Apparatus, MA, US), as in previous studies from our lab [36–39]. Horizontal activity is reported as both total distance traveled during the entire session, as well as in separate 5-min bins that allow for the detailed investigation of the effects of PLN ablation on locomotor activity [36, 39, 40].

#### 2.5.2. Y-maze Test

Spatial working memory was assessed using the Y-maze task, as previously described [41, 42]. Briefly, mice of both sexes were individually placed in the center of a Y-shaped fiber plastic light- colored opaque maze comprised of three identical arms (A, B and C), measuring 22cm x 6.4cm x 15cm (120° from each other), and were allowed to freely explore all three arms over a 10-min period. A CCD camera fixed above the center of the maze allowed for digital video output and performance in the Y-maze task was manually scored. Spontaneous alternation (per cent) was measured as an index of spatial working memory driven by the animal’s innate preference to explore novel areas [43].

Spontaneous alternations (SA) were defined as successful entries to three separate arms consecutively. Subsequently the SA% score was calculated using the formula below:

SA% = [Number of spontaneous alternations/(Total number of arm entries-2)] x 100

#### 2.5.3. Object Recognition and Locational Memory

Long-term object recognition memory (ORM) and object locational memory (OLM) were assessed using a two part protocol, as previously described [44]. The same mouse cohort first completed the OLM testing, followed by the ORM testing with 7 days (d) apart. Briefly, mice were subjected to 3 consecutive phases: i) habituation (6d), ii) training (1d), and iii) testing (1d). All phases were conducted at dim light conditions (∼47 Lux). During habituation, each mouse was introduced to an open non- translucent arena measuring 45 cm x 45 cm x 45 cm for 5 min per day and for 6 consecutive days. During the OLM training, two identical objects were placed in the arena in symmetrical positions. Each mouse was allowed to freely explore the arena for 10 min. The OLM test was conducted at 24h following the training phase, and one object was moved to the opposite side of the arena. Mice were allowed to explore the arena for 5 min. At 7d following completion of OLM testing, mice were habituated to the circular ORM arena (24 cm in diameter). In the ORM training, two identical objects were placed into the arena in symmetrical locations and mice were allowed to explore the arena for 10 min. Two sets of objects were used in the ORM test to avoid innate preference for one object; familiar and novel objects were counterbalanced across groups. In the ORM testing, one object was replaced with a novel object of different shape and color, and mice were allowed to explore the arena for 5 min. All sessions were video-recorded and blindly scored for time spent interacting with each object.

#### 2.5.4. Three-chamber Social Interaction Test

Sociability and social memory were assessed as previously described [45, 46]. Briefly, mice were individually introduced to a partitioned box and upon habituation underwent a two-session social test protocol. The first session assessed sociability, while the second session tested for preference for social novelty, that is indicative of social memory, as in our previous study [46]. The social test apparatus consisted of a transparent acrylic box with removable partitions, divided into three chambers; two equally sized distal chambers (20cm x 40cm, each) and one middle chamber (20cm x 17.5cm), with 5 cm openings between each chamber. Two cylindrical wire cages (10 cm in height, 9 cm bottom diameter) were used to contain the social stimuli. For the sociability test, the test animal was habituated to the middle chamber for 5 min. Subsequently, an unfamiliar wild-type mouse of the same sex (*i.e.*, Stranger I) was positioned under one of the wire cages located in one of the distal chambers, while an empty cage was inserted in the other distal chamber. The dividers were then raised, and the test animal was allowed to freely explore all three chambers for 10 min. Following this session, a second unfamiliar mouse of the same sex (*i.e.*, Stranger II) was inserted into the previously empty wire cage, and the test animal was left to continue exploring all chambers for 10 min. Time spent in close interaction with both cages was recorded. The release of the animals and the relative positioning sequence of the stranger mice were counterbalanced across different animals but were kept consistent for each individual test animal. All sessions were video-recorded and blindly scored for time spent interacting with each object.

#### 2.5.5. EEG-based Analysis of Sleep Architecture

Sleep architecture was assessed using electroencephalogram (EEG)/electromyogram (EMG)-based polysomnography, as previously described [46–48]. Briefly, anesthetized mice underwent aseptic stereotaxic surgery for the implantation of epidural EEG electrodes in the format of a commercially available prefabricated head mount (Pinnacle Technology, Lawrence, KS). The headmount consisted of a plastic 6-pin connector glued to a printed circuit board (PCB), where three EEG electrodes and two EMG electrodes were affixed. Four stainless-steel screws secured the head mount on the skull and served as the EEG electrodes. The two anterior screws (0.1 inches length) were placed 1.5 mm lateral to the midline and 1 mm anterior to the bregma, while the two parietal screws (0.125 inches length) were placed 1.5 mm lateral to the midline and approximately 2 mm anterior to lambda. Conductivity with the PCB was secured by silver epoxy application. In addition, two 1.5 cm long stainless-steel wires served as EMG electrodes and were embedded in the neck musculature. Upon fixation of the structure on the skull with dental cement, the neck incision was closed with sterile non- absorbing acrylic sutures and a topical analgesic was applied. Buprenorphine analgesia was provided, as needed. The mice were then left to recover and habituate in the circular recording cages for 7-10 days prior to being connected to the recording cable (*i.e.,* pre-amplifier). Two days later, a polygraphic recording of the spontaneous sleep–wakefulness states was performed during 48 h, beginning at 7:00 pm. Subsequently, EEG/EMG recordings were blindly scored in 10 sec epochs as wakefulness, non- rapid eye movement (NREM) sleep, or rapid eye movement (REM) sleep, following classical criteria [49, 50] using the Sirenia® analysis package (Pinnacle Technology, Inc. KS, USA). Spontaneous sleep–wakefulness patterns and the amounts of vigilance states were determined for each mouse throughout 48h and summed over 24h, as in our previous studies [46]. For bout analysis, a bout was defined as 3 or more consecutive epochs of one vigilance state. Total bout number and average bout length for wakefulness, NREM, and REM sleep were reported for the total recording period, dark period, and light period. Arousals during NREM sleep were defined as 1-2 epochs (i.e. below the bout threshold) of wakefulness during a period of NREM sleep. Sleep spindle analysis was not conducted as it is not a feature of the Sirenia® analysis package. Spectral analysis was conducted using the Fast Fourier Transform (Sirenia Sleep Pro, Pinnacle Technology Inc) to calculate the power (μV2) of delta (δ; 0.5-4Hz), theta (θ; 4-8Hz), alpha (α; 8-13Hz), beta (β; 13-30Hz), and gamma (γ; 30-40Hz), frequency bands during each of the sleep stages. Total power of each sleep state was summed and normalized as a percentage of total power across all frequency bands.

#### 2.5.6. 5-Choice Serial Reaction Time Task (5-CSRTT) for the Assessment of Executive Functioning

Impulsivity and attention were assessed using the 5-CSRTT in a 9-hole operant chamber using protocols provided by Lafayette Instrument Co (Lafayette Instrument Co., IN, USA) and based on previous studies from our lab and others [51, 52, 74]. Male and female control and cKO were single housed throughout a reverse 24/h light/dark cycle, and all testing was performed during dark hours. Prior to training, mice were food restricted to 90% of free-feeding body weight. Following food restriction, each mouse was pre-exposed to 25mL of the reward (Yoohoo; Mott’s, Plano, TX, USA) at 24 hours before magazine training to reduce neophobic reactions to the reward. The 5CSRTT chamber features on one side a reward tray where liquid reward is dispensed through a pump, and on the other side 9 nose-poke holes with recessed lights. Nose-pokes were detected via infrared beam break and recorded automatically. Each hole and the reward tray have a light that illuminates conditionally. For this experiment, only nose-pokes in holes 1, 3, 5, 7, and 9 were recorded. Holes 2, 4, 6, and 8 remained inactive throughout all 5-CSRTT training and testing. Following pre-exposure to reward, all mice began the magazine training schedule. During magazine training, the center nose-poke hole illuminated for 8 sec, followed by dispensal of reward solution. Upon collecting the reward solution, a 5 sec inter-trial interval (ITI) occurred prior to the next trial. Mice were required to complete 30 trials within the 30 min session to progress to the next phase. After magazine training, all mice were moved into a fixed ratio schedule (FR1). In this schedule, the center nose-poke hole remained lit until a nose-poke response was made, at which point it terminated and reward was dispensed. A magazine entry was required for the next trial to start. Mice completed a single session (30 min/50 trials, whichever was reached first) each day for 17 days (d). Criterion was 50 completed trials; mice that reached criterion within 17d of FR1 training were passed onto a progressive ratio (PR) session. Following the FR1, all mice completed a single PR session. This session was identical to the FR1, except that the number of nose-pokes required to dispense reward increased every third trial (i.e., 1,1,1,2,2,2,4,4,4,7,7,7…). The PR session was conducted for 60 min, or until no response was made for 5 min, defined as the breakpoint. After completion of the PR session, mice progressed to the basic 5-CSRTT testing paradigm. In this paradigm, a session begins with a dispensal of reward. After entry into the reward magazine, a 5 sec ITI is initiated, during which all lights are off and the animal is required to inhibit its response. After the ITI, one of the five odd-numbered holes illuminated for a duration that remained constant for that session, followed by a 3 sec limited hold where the light turned off but responses were still accepted. A nose-poke to the illuminated hole was recorded as a correct response, whereas a nose-poke to another hole was recorded as an incorrect response. If no response was made to any hole during the stimulus duration or the 3 sec limited hold, an omission was recorded. Finally, nose-pokes during the ITI were recorded as premature responses. Reward was dispensed only after correct responses were made. All animals completed a single 30 min session each day, and performance measures were reported for the first day criteria was met (>20 responses, with 50% response accuracy). Basic 5-CSRTT experiments were completed in 3 sub-phases with progressively decreasing stimulus duration (32 sec, 16 sec, 8 sec). In order to progress in these sub-phases, mice needed to reach criteria for two consecutive sessions. Mice progressed at their own pace and were subjected to a single titration 5-CSRTT session following completion of basic 5-CSRTT testing. In the titration session, mice were subjected to a more difficult version of the basic 5-CSRTT. In this session, stimulus duration was titrated based upon an animal’s performance within that session. The stimulus duration was set at 3 seconds for the first 3 trials; upon a correct response, the stimulus duration for the following trial was shortened one step in the series down to a minimum of 0.1 s, whereas an incorrect response or omission lengthened the stimulus duration one step. Premature responses did not change the stimulus duration on the subsequent trial. A limited hold was present such that the animal always had 3 sec to make a response (i.e., 0.5 sec stimulus duration + 2.5 sec limited hold). The ITI remained constant (5 sec) for all trials. Each animal completed a single 30 min session of the titration 5-CSRTT. For each session, the following behavioral endpoints were calculated and defined as follows: i) Accuracy (i.e., correct responses/total trials); ii) Omissions (i.e., omitted trials/total trials); iii) Correct Response Latency (i.e., latency to make a correct response following stimulus presentation); iv) Response Accuracy (i.e., correct responses/correct+incorrect responses) and v) Premature Response Rate (i.e., premature responses/correct+incorrect+premature responses).

### 2.6. Statistical Analysis

All data are presented as means±SEM. Data were analyzed using two-way analysis of variance (ANOVA) with genotype and sex as independent factors, followed by appropriate *Bonferroni post-hoc* analysis in view of statistically significant interactions. In cases where the *Levene’s* test of homogeneity was significant the effects of genotype were assessed using non-parametric *Kruskal-Wallis* tests. Temporal locomotor and sleep data were analyzed using three-way repeated measures (RM) ANOVA (i.e., factors: genotype, sex and time). Homogeneity of covariance was assessed using the *Mauchly’s* test of sphericity; in cases where sphericity was violated the RM ANOVA F was modified according to the *Greenhouse-Geisser* correction to make it more conservative. Performance in the FR1 Session of the 5-CSRTT was analyzed using Kaplan-Maier survival analysis followed by Mantel-Cox Log Rank pairwise comparisons. The IBM SPSS Statistics for Windows, Version 24.0 (Armonk, NY: IBM Corp.) and the GraphPad Prism version 8.0.0 for Windows (GraphPad Software, La Jolla, CA USA) were used for statistical analysis and graph plotting. Statistical significance was defined as p ≤ 0.05. Detailed statistical analyses are reported in **Supplementary Table S1.**

## 3. RESULTS

### 3.1. PLN is Expressed in the GABAergic TRN Neurons

In a first set of experiments, we investigated the expression of PLN in the mouse brain **(Fig. 2).** We found *Pln* mRNA to be expressed at relatively low levels in the adult mouse brain using RT-PCR **(Fig. 2a).** Follow-up IHC analysis showed PLN protein expression to be confined in the GABAergic GAD67+ neurons of the TRN **(Fig. 2bc)** and for PLN to colocalize with SERCA2 in the adult mouse TRN **(Fig. 2d).** In our experimental setup we did not detect PLN immunoreactivity in other brain regions, including the striatum (STR), thalamus (TH), prefrontal cortex (PFC), hippocampus (HIPP), and cerebellum (CER) **(Fig. 3),** or in other cell types (e.g., astrocytes; **Supplementary Fig. S1**) indicating that PLN may possibly play a unique role in the GABAergic TRN neurons of the mouse brain.

**Figure 2:**
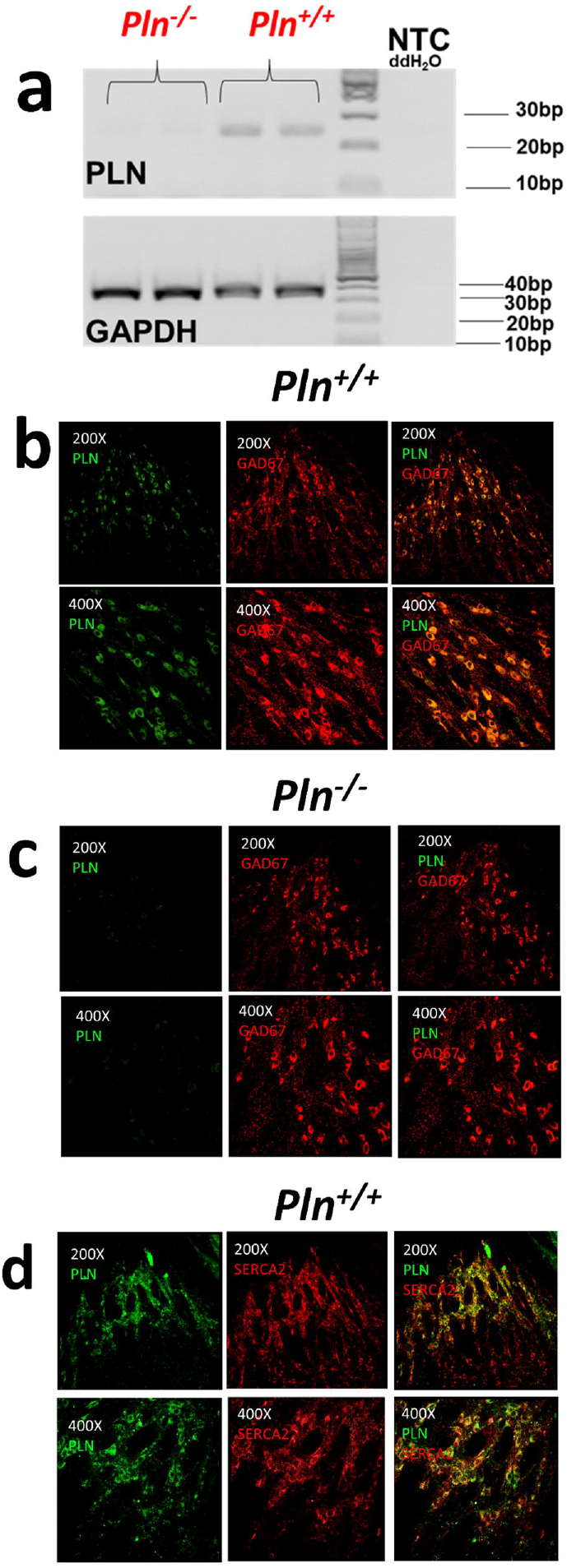
PLN is expressed in the TRN of the mouse brain: **a)** PLN mRNA was detected in the whole brain of adult *Pln^+/+^* mice but not in *Pln^-/-^* mice. PLN-specific immunoreactivity is detected in the GABAergic GAD67^+^ TRN neurons of the adult mouse TRN in **b)** *Pln^+/+^* but not in **c)** *Pln^-/-^* mice; **d)** PLN colocalizes with SERCA2 in the TRN neurons of *Pln^+/+^* mice. PLN, GAD67 and SERCA immunoreactivity was assessed using fluorescent IHC in coronal free-floating brain sections from adult *Pln^+/+^*and *Pln^-/-^* mice stained with mouse monoclonal PLN (1:100), rabbit polyclonal GAD67 (1:250) and rabbit polyclonal SERCA2 (1:500) primary antibodies. Representative images shown from at least three independent runs were obtained at 200X and 400X magnifications using an Olympus Fluoview 3000 confocal microscope (Olympus, MA, USA).

**Figure 3:**
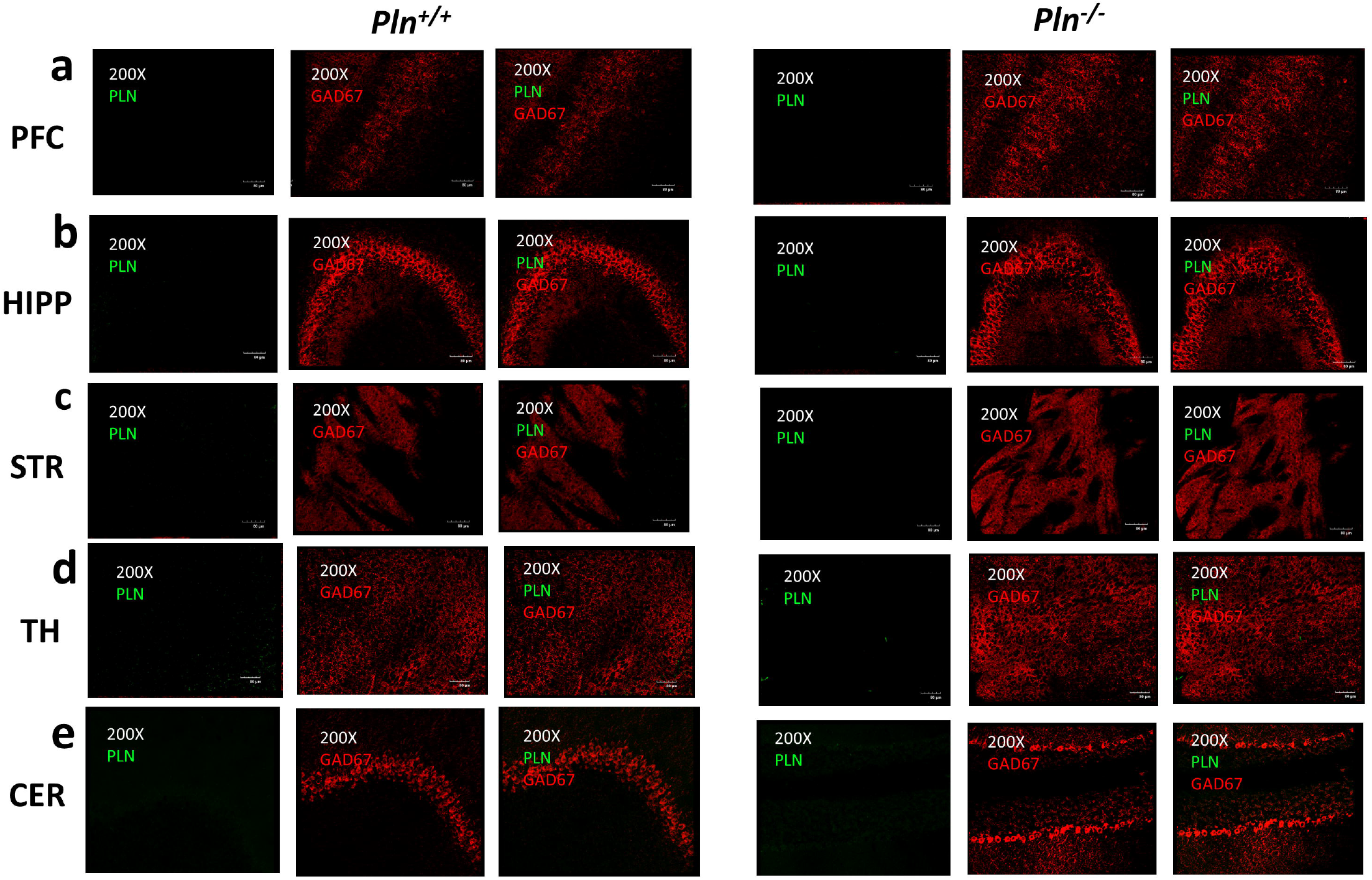
PLN immunoreactivity is not detected in other brain regions: PLN immunoreactivity was not detected in other brain regions, including: **a)** the prefrontal cortex (PFC); **b)** the hippocampus (HIPP); **c)** the striatum (STR); **d)** the thalamus (TH) and **e)** the cerebellum (CER). PLN and GAD67 protein expression was assessed using fluorescent IHC in coronal free-floating brain sections from adult *Pln^+/+^*and *Pln^-/-^* mice stained with mouse monoclonal PLN (1:100) and rabbit polyclonal GAD67 (1:250) primary antibodies. Representative images were obtained at 200X and 400X magnifications using an Olympus Fluoview 3000 confocal microscope (Olympus, MA, USA).

### 3.2. Constitutive Global Loss of PLN Induces Behavioral Deficits in Mice

Upon confirming that PLN is expressed at the protein level in the TRN, we employed an existing global *Pln^-/-^* knockout mouse model to assess whether genetic ablation of PLN induces major behavioral effects at the organismal level. Locomotor analysis in the OFT revealed that *Pln^-/-^*mice were hyperactive, as compared to their *Pln^+/+^* littermates **(Fig. 4a),** and spent significantly more time traveling in the center of the OFT arena, indicative of anxiolysis **(Fig. 4b)**. Moreover, genetic *Pln* ablation affected mouse performance in the Y-maze test indicating that *Pln^-/-^* mice present a deficit in spatial working memory **(Fig. 4c)**. Even though global *Pln* deletion did not significantly affect object location memory (OLM), as both *Pln^-/-^ and Pln^+/+^* littermates spent significantly more % time interacting with the displaced object during the testing phase of the OLM test **(Fig. 4d)**, it significantly impacted object recognition memory (ORM), as *Pln^-/-^* mice spent almost equal time exploring the novel and the familiar objects during the testing phase of the ORM **(Fig. 4e)**. Analysis of social exploratory behavior showed that global *Pln* deletion did not significantly affect sociability **(Fig. 4f)** and preference for social novelty **(Fig. 4g)** indicating that *Pln* ablation does not significantly affect sociability in mice. The results of the statistical analyses are reported in **Supplementary Table S1.**

**Figure 4:**
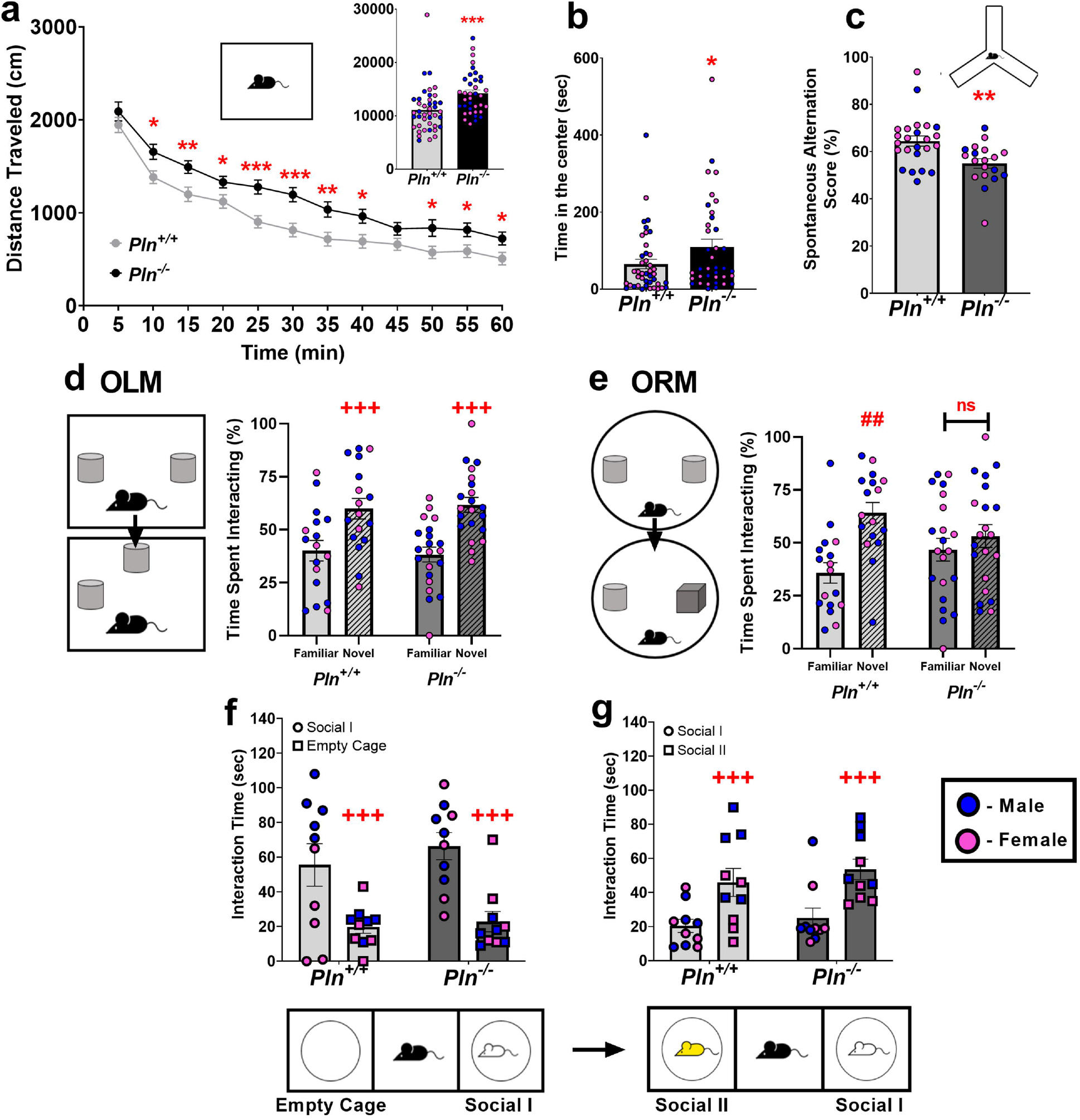
Global genetic ablation of *Pln* induces hyperactivity and cognitive deficits in mice: Locomotor analysis in the Open Field test (OFT) revealed that *Pln^-/-^* mice are hyperactive **(a)** and spend significantly more time in the center of the OF arena **(b)**, as compared to their *Pln^+/+^* littermates; **c)** performance in the Y-maze is impaired in *Pln^-/-^* mice; **d)** Object location memory (OLM) was not affected in *Pln^-/-^* mice; but **e)** global loss of PLN function was associated with impaired object recognition memory as *Pln^-/-^* mice spent the same time exploring the novel and the familiar objects in the ORM test; **f)** sociability and **g)** preference for social novelty was not affected by global PLN deletion. Individual values for male and female mice for each experiment are shown; *p<0.05; **p<0.01; ***p<0.001: statistically significant differences as compared *to Pln^+/+^* mice; ^+++^p<0.001 main effect of object or stimulus as analyzed by 2-way ANOVA; ^##^p<0.01: statistically significant difference between familiar and novel object in *Pln^+/+^* mice.

### 3.3. Conditional *Pln* Deletion in TRN Neurons Results in Hyperactivity

To further explore the TRN-dependent functions of PLN, we generated **(Fig. 1)** and validated a novel Cre-LoxP conditional knockout (cKO) mouse model with a targeted deletion of *Pln* in GABAergic TRN neurons **(Fig. 5)**. Given that our immunohistochemical data revealed that expression of the PLN protein in the mouse brain is restricted to GABAergic neurons of the TRN, we selected a well-defined GABAergic marker of TRN neurons (i.e., GAD2), as our Cre driver mouse line in order to selectively ablate PLN protein expression in TRN neurons while leaving expression in other tissues such as cardiac, skeletal, and smooth muscle intact [18, 21]. Using double fluorescent IHC we confirmed that the *Pln* gene is effectively deleted in the brain’s TRN, as assessed by the absence of PLN immunoreactivity in the GAD67^+^ TRN neurons in cKO mice **(Fig. 5a)**, while PLN protein is expressed peripherally in the cardiac muscle of cKO mice **(Fig. 1f).** Notably, analysis of locomotor activity in the OFT showed that the conditional deletion of *Pln* in the TRN induces hyperactivity in cKO mice **(Fig. 5b,c),** further confirming our findings in *Pln^-/-^* mice and consolidating a role for PLN in regulating locomotor activity. Mouse performance in the Y-maze was not affected in cKO mice suggesting that unlike global *Pln* deletion, TRN-specific loss of *Pln* function does not impact spatial working memory **(Fig. 5d)**. The results of the statistical analyses are reported in **Supplementary Table S1.**

**Figure 5:**
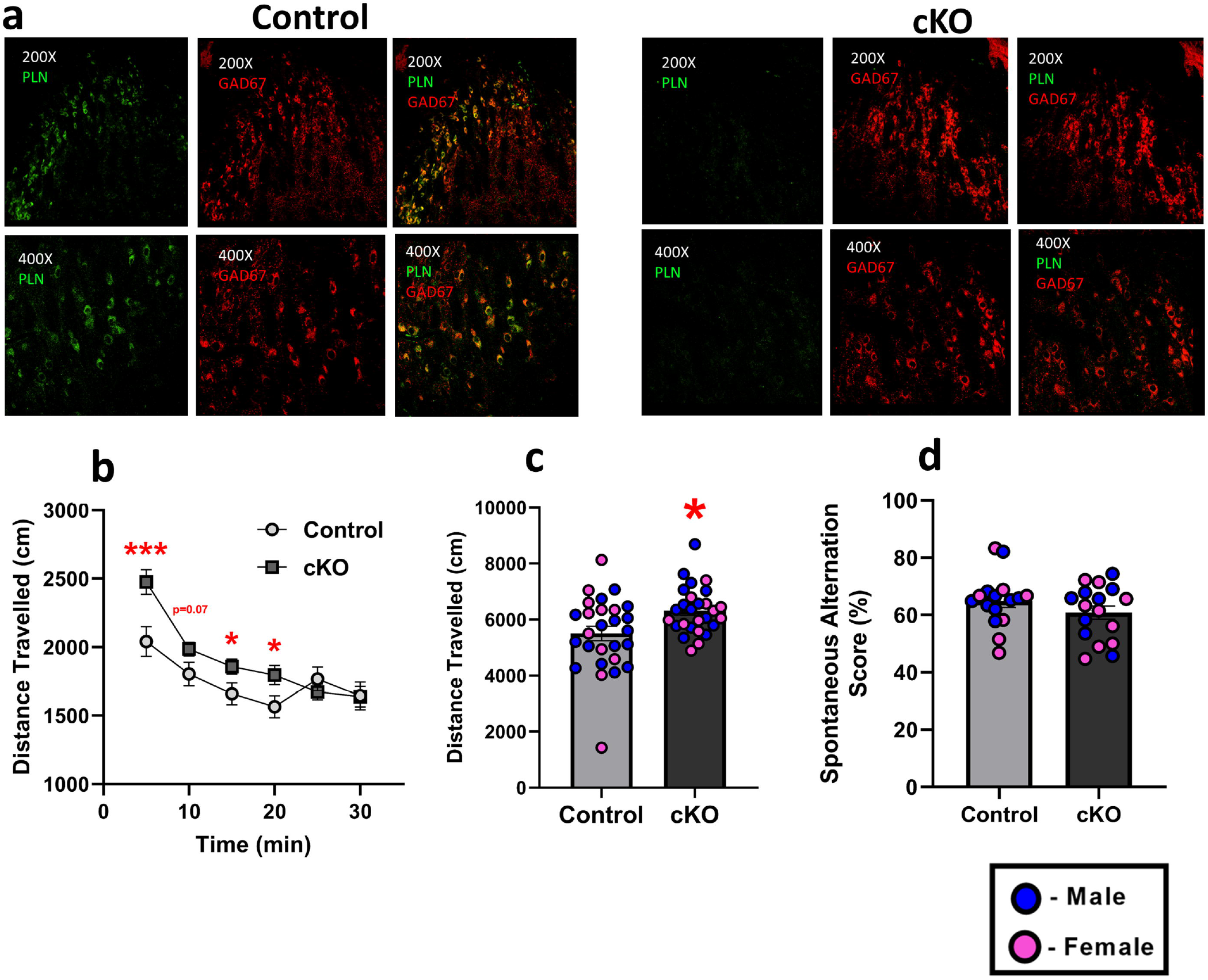
Generation of the floxed PLN mouse line and validation of the conditional knockout (cKO) mice. **a)** Fluorescent IHC confirmed the conditional deletion of *Pln* in the GABAergic TRN neurons as PLN immunoreactivity was detected in the TRN neurons of control, but not in their cKO littermates; representative images shown from at least three independent runs were obtained at 200X and 800X magnifications using an Olympus Fluoview 3000 confocal microscope (Olympus, MA, USA). Conditional deletion of PLN in the TRN induces **b)** a hyperactive phenotype in cKO mice with **c)** cKO mice traveling a greater distance in the OF during the first 15 min of the test; **d)** spatial working memory was not affected in cKO mice, as assessed in the Y-maze. Individual values for male and female mice for each experiment are shown; *p<0.05; ***p<0.001: statistically significant differences between control and cKO mice.

### 3.4. Loss of PLN function in the TRN Affects Sleep Architecture and Impulsivity in cKO mice

EEG/EMG polysomnography for the analysis of sleep architecture revealed that conditional deletion of *Pln* in the TRN resulted in pronounced alterations in sleep architecture that were confined in the dark period of the cycle **(Fig. 6).** Specifically, cKO mice spent less time in wakefulness **(Fig. 6d)** and greater time in NREM **(Fig. 6e)** and REM sleep **(Fig. 6f)** during the dark period, as compared to their control littermates. Moreover, cKO mice also had shorter wakefulness bouts in the dark period **(Fig. 6d)**, a higher number of NREM bouts in the dark period **(Fig 6e)**, and longer REM bouts in the dark period as compared to the control mice **(Fig. 6f).** cKO mice also exhibited increased transitions between wakefulness and NREM, **(Fig. 6g)** and a trend towards increased incidence of arousals during NREM sleep **(Fig. 6i)** during the dark period. The same analysis for REM sleep in the dark period revealed that cKO mice spent more time in REM sleep as compared to their control counterparts; *post-hoc* analysis indicated that the prolonged REM sleep duration during the dark period was more pronounced in female mice. EEG spectral analysis did not reveal any statistically significant differences between the spectral profiles of control and cKO mice in any power band or vigilance state in the dark or light period **(Supplementary Table S2)**. Behavioral analysis in the 5- CSRTT indicated that conditional loss of PLN function in the TRN affects executive functioning and results in impulsivity. Specifically, in the FR1 session, a Kaplan-Maier survival analysis followed by Mantel-Cox Log Rank pairwise comparisons revealed that female cKO mice presented a delay in reaching the FR1 criterion as compared to the other genotypes indicating that conditional deletion of *Pln* in the TRN affects the initial learning acquisition rate of female mice during FR1 testing of the 5- CSRTT; however, despite this initial deficit, all mice eventually acquired the task **(Fig. 7ab)**. No statistically significant differences were observed in the PR **(Fig. 7c)** and the main 5-CSRTT sessions *(data not shown)*. Most importantly, in the final and more challenging “Titration” 5-CSRTT session, attention was not affected in cKO mice **(Fig. 7d-i),** but overall cKO mice displayed an impulsive phenotype as evidenced by a trend for increased premature response rate **(Fig. 7j)** and increased number of premature responses **(Fig. 7k).** The results of the statistical analyses are reported in **Supplementary Table S1.**

**Figure 6:**
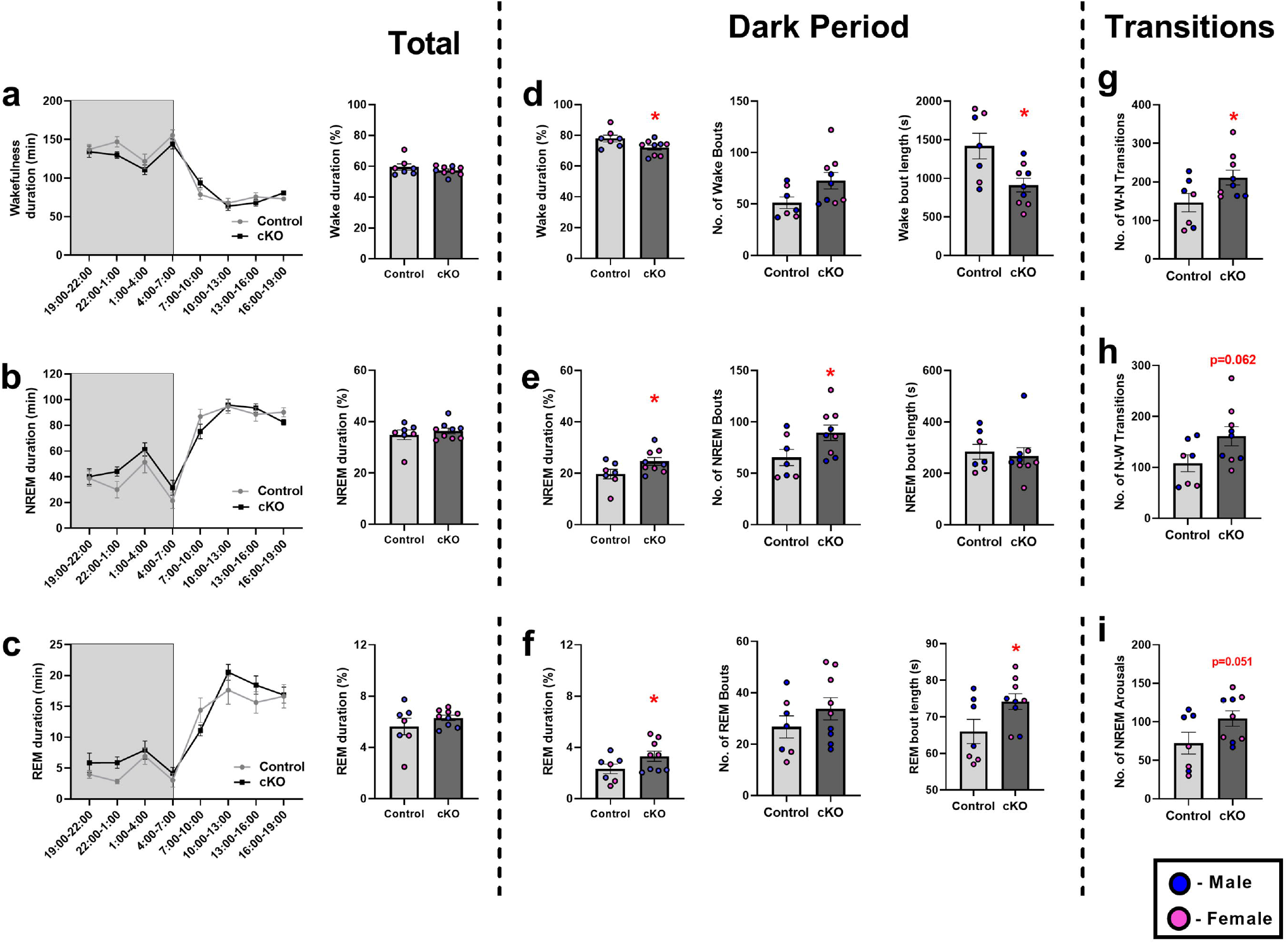
Conditional deletion of *Pln* in the TRN affects sleep architecture. Analysis of EEG/EMG polysomnographic recordings indicate that **a-c)** both cKO and control mice exhibit the typical polyphasic structure of vigilance states found in rodents and a diurnal rhythm of sleep and wakefulness, with larger amounts of sleep during the light period classically observed in nocturnal species. **d-f)** Interestingly, cKO mice exhibit significantly increased amounts of NREM and REM sleep at the expense of wakefulness during the dark period, as well reduced wakefulness bout length in the dark period, increased NREM bouts in the dark period, and increased REM bout length in the dark period. **g-i)** cKO mice also exhibit enhanced dark period transitions between wakefulness and NREM, and a trend towards increased arousals during NREM in the dark period. Individual values for male and female mice for each experiment are shown; *p<0.05 statistically significant difference between control and cKO mice.

**Figure 7:**
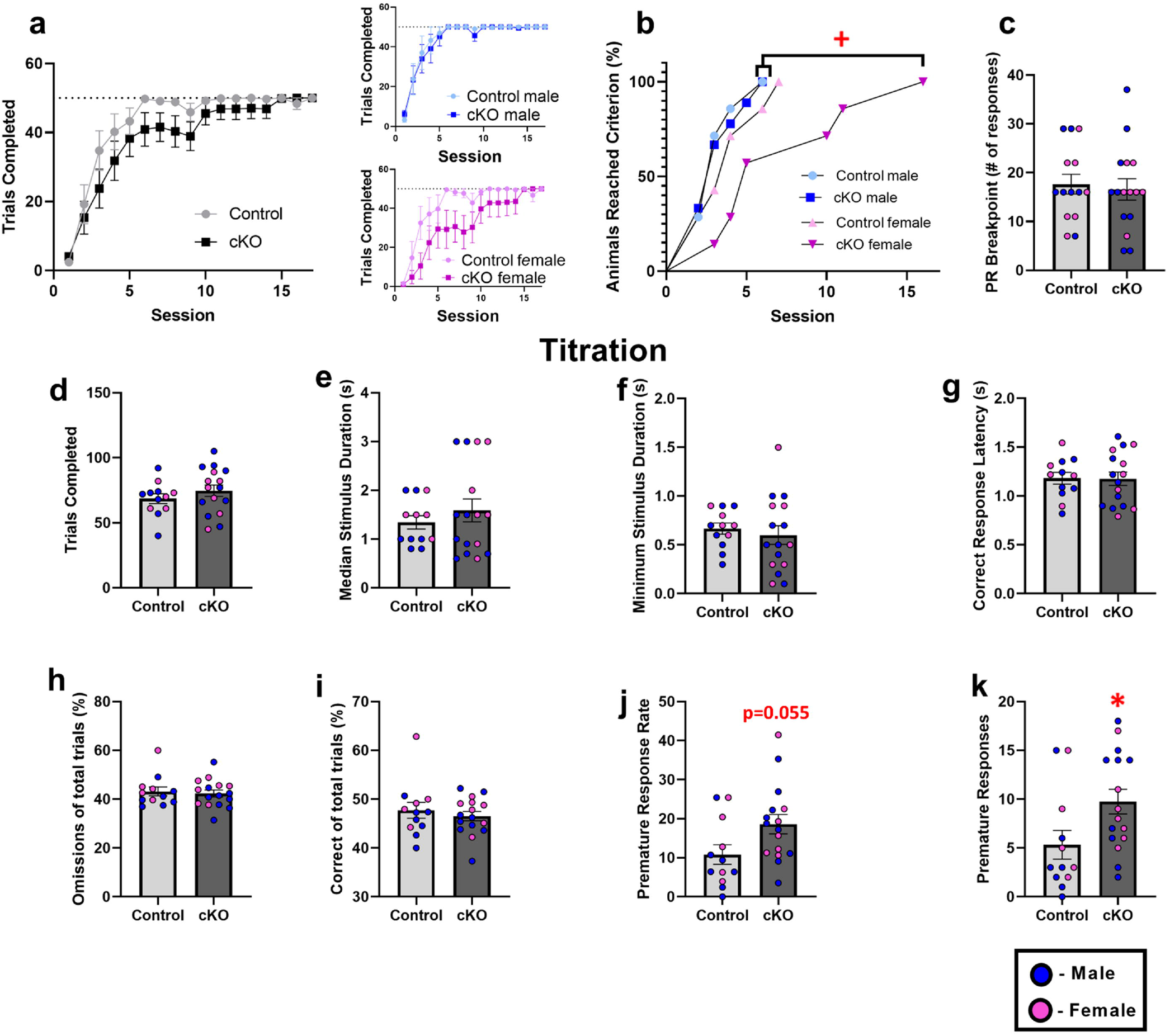
Effects of conditional loss of PLN function in the TRN on executive functioning. **a)** Trials completed each day of the FR1 task in control and cKO mice of both sexes. **b)** Survival curve analysis revealed that female cKO mice exhibit a significant delay in the acquisition of the FR1 task, as compared to control and cKO male mice, as well as a statistical trend as compared to female control mice *(p=0.075)*. **c)** cKO mice do not exhibit deficits in the PR task. **d-i)** Conditional deletion of PLN in the TRN did not significantly impact attentional processes as assessed by number of total trials, % correct responses, % omitted responses, correct response latency, median stimulus duration, or minimum stimulus duration in the final “Titration” session of the 5-CSRTT; **j)** In the 5-CSRTT titration task, cKO mice displayed an impulsive behavior evidenced by an enhanced premature response rate and **k)** number of premature responses.*p<0.05: statistically significant differences between control and cKO mice; ^+^p<0.05 statistically significant differences between cKO females and control males and cKO males.

## 4. DISCUSSION

This is the first study to investigate the role of PLN in brain and behavior. Herein, we report that PLN is expressed in the TRN of the adult mouse brain and that loss of PLN function results in robust behavioral deficits, including hyperactivity, impulsivity and alterations sleep architecture in mice.

Despite the fact that the SERCA2 is ubiquitously expressed in neurons [8, 28, 29], earlier Northern blotting studies had reported that the *Pln* gene is not expressed at the mRNA level in the brain of humans, pigs, or mice [30, 31, 53]. More recently, a number of high-throughput global *RNA-seq* and genome-wide microarray gene expression profiling studies have detected *Pln* mRNA in individual mouse dorsal root ganglion (DRG) neurons, immunopanning-isolated mouse and human cortical neural cells, including astrocytes and neurons, as well as in fetal human primary astrocytes [54–57], supporting the expression of PLN in neural tissues. In our experimental setup, low *Pln* mRNA levels were detected in the whole mouse brain **(Fig. 2a),** whereas detection of PLN protein expression was exclusively confined in the GABAergic neurons of the TRN in wild-type *Pln*^+/+^ mice **(Fig. 2b)**, but not in their *Pln*^-/-^ knockout counterparts **(Fig. 2c)**. As theoretically expected, we also confirmed that PLN protein expression co-localizes with SERCA2 in the mouse TRN neurons **(Fig. 2d)**. Importantly, PLN immunoreactivity was exclusively observed in GABAergic TRN neurons but not in other brain regions studied (i.e., the striatum, the prefrontal cortex, the thalamus, the hippocampus, and the cerebellum) **(Fig. 3)** or in other cell types, (i.e., astrocytes**; Fig. S1).** Of note, our fluorescent immunohistochemistry data corroborate recent bulk *RNA-seq* studies reporting higher *Pln* mRNA expression in the mouse thalamus [58], and the TRN-specific PLN mRNA expression pattern observed by *in situ* hybridization histochemistry (ISH) in the Allen Mouse Brain Atlas [e.g., Exp: 69116799 (sagittal) & Exp: 74357563 (coronal)] [59]. Taken together, our findings indicate that PLN protein is selectively expressed in the GABAergic TRN neurons and provide first evidence that the PLN/SERCA2 pathway operates in the central nervous system where it possibly plays a unique role in regulating intracellular Ca^2+^-handling in the TRN neurocircuitry.

TRN dysfunction has been proposed to underlie hyperactivity, deficits in executive functioning, and sleep disturbances [5] and to comprise a circuit endophenotype in neurodevelopmental disorders, including autism spectrum disorder (ASD), schizophrenia and attention-deficit/hyperactivity disorder (ADHD) (for review see [60]). Thus, selective expression of PLN in the TRN further suggests that the PLN/SERCA2 pathway is of critical importance in regulating the dynamics of the TRN neurocircuitry and fundamental behavioral processes of great biological significance. Indeed, compelling evidence suggests that SERCA plays a critical role in brain physiology and pathophysiology; SERCA2 dysregulation has been associated with several neuropsychiatric disorders, including schizophrenia, bipolar disorder, cerebral ischemia, alcoholism, as well as Alzheimer’s and Parkinson’s diseases (for review see [17, 61]). Interestingly, recent studies from our group have revealed that chronic pharmacological stimulation of SERCA induces anxiety- and depressive-like behaviors, memory deficits, as well as sustained brain region-specific noradrenergic and serotonergic neurochemical alterations in mice, further highlighting the importance of dissecting the neurobiological implications of SERCA2-dependent Ca^2+^ regulatory mechanisms in the brain [39, 74].

Interestingly, TRN-specific dysfunction has been linked to an ADHD-like hyperactive phenotype in mice as TRN-targeted genetic ablation of the patched domain containing 1 (*Ptchd1*) gene results in locomotor hyperactivity [62]. Thus, upon confirming the expression of PLN in the mouse TRN, we further sought to assess the behavioral consequences of global genetic ablation of *Pln* using an existing *Pln* knockout mouse model [32, 33]. Interestingly, constitutive deletion of the *Pln* gene resulted in hyperactivity and cognitive deficits in *Pln^-/-^* mice as they traveled a greater distance in the OFT **(Fig. 4a),** and were outperformed by their *Pln^+/+^*counterparts in the Y-maze and the NOR tests, with these behavioral deficits being indicative of impaired spatial working memory and long-term object recognition memory, respectively **(Fig. 4c-e)**. Interestingly, the behavioral deficits observed in *Pln^-/-^* mice were not accompanied by social deficits as assessed in the three-compartmental sociability test **(Fig. 4fg),** supporting the notion that PLN is not strongly implicated in the behavioral ASD-relevant domain.

It is well established that the TRN regulates important brain-wide functions, including sensorimotor processes [64], executive functioning [62] and the generation of sleep rhythms [63]. To further explore the TRN-dependent functions of PLN, we generated and validated a novel conditional cKO mouse line with diminished PLN expression in the GABAergic TRN neurons. Indeed, we found that conditional deletion of *Pln* in the TRN disturbed thalamic reticular-dependent behaviors.

Specifically, much like their *Pln^-/-^* 129Sv counterparts, cKO C57 mice were hyperactive **(Fig. 5bc)**, while deletion of the *Pln* gene in TRN neurons was associated with impulsivity, as observed in the titration session of the 5-CSRTT **(Fig. 7j,k)**. As no differences were observed in reward collection latency *(data not shown)* nor correct response latency **(Fig. 7g)**, the hyperactive phenotype of cKO mice does not significantly confound the interpretation of the 5-CSRTT data [65]. Despite the fact that cKO mice did not exhibit deficits in spatial working memory in the Y-maze, female cKO mice presented a delay in reaching the FR1 criterion in the 5-CSRTT **(Fig. 7ab)** indicating that conditional loss of PLN function in the TRN may result in an initial sex-specific associative learning deficit in acquiring this operant conditioning task, without this deficit however significantly affecting attentional processing in the subsequent phases of the 5-CSRTT [51]. Notably, the TRN is critical for the generation of sleep spindles and NREM sleep [63, 66, 67]. Thus, it is conceivable that deregulation of SERCA2-dependent Ca^2+^ signaling mechanisms may impact thalamocortical network oscillations relevant for NREM sleep and overall sleep architecture. Present data revealed that cKO mice exhibited significantly increased amounts of NREM and REM sleep, at the expense of wakefulness, throughout the dark period **(Fig. 6d,e,f).** In line with these findings, bout analysis of the dark period revealed cKO mice exhibited more NREM and REM bouts, enhanced REM bout length, and reduced wakefulness bout length **(Fig. 6f)**. Interestingly, previous studies have reported fragmented sleep architecture in mice with ADHD-like behavioral features [62], while genetic ablation of CaV2.3 Ca^2+^ channels, that are predominantly expressed in the TRN, has been reported to result in increased NREM sleep duration at the expense of wakefulness in CaV2.3KO mice [68]. As TRN neuronal firing patterns have been shown to directly mediate the transition between wakefulness and NREM [69], we hypothesize that a potential mechanism for these sleep abnormalities in cKO mice may include aberrations in Ca^2+^-mediated TRN neuronal firing due to loss of PLN, enhancing wakefulness-NREM transition-promoting TRN circuits.

It is noteworthy that differences between the behavioral profiles observed between global 129S *Pln^-/-^* and conditional C57 cKO mice could be attributed to the putative peripheral effects of PLN, and/or to the different mouse strains used. Notably, however, locomotor hyperactivity upon loss of PLN function was a robust behavioral finding that was evidenced upon both global and conditional deletion of *Pln* and across both mouse strains underscoring the role of PLN in regulating locomotor activity though the thalamic reticular neurocircuit. Interestingly, a novel subcortical GABAergic neural input pathway was recently discovered that originates in the TRN and projects to striatal parvalbumin (PV) interneurons [70]. Theoretically, excitation of this TRN projection would result in inhibition of striatal PV interneurons and concomitant disinhibition of the striatal spiny projection neurons (SPNs) [70] that comprise as much as 95% of the total striatal neuron population [71], thus providing a putative subcortical pathway by which the TRN could directly affect striatum-dependent behaviors, such as locomotor activity and executive functioning. Interestingly, in our experimental setup the core behavioral consequences of genetic *Pln* ablation were not sex-differentiated; however, two peripheral behavioral measures (i.e., time to reach FR1 criterion in the 5-CSRTT and REM sleep) appeared to be differentially affected between male and female cKO mice underscoring the need for future studies to employ both sexes when assessing the effects of PLN in the brain. Studies from our lab are underway to interrogate the role of the PLN/SERCA2 pathway in specific components of the TRN neurocircuitry, to investigate the effect of *Pln* ablation on thalamic reticular Ca^2+^ dynamics and electrophysiological neuronal properties in order to dissect the cellular and molecular mechanisms pertaining to the role of PLN in the TRN.

## 5. CONCLUSIONS

Overall, our study shows that loss of PLN function in the TRN results in aberrant thalamic reticular behavioral phenotypes in mice (i.e., hyperactivity, impulsivity and sleep deficits). Present data exposed a novel role for PLN in the brain, and further suggest that PLN is a critical regulator of SERCA2 in the TRN neurocircuitry; the intricate molecular mechanisms orchestrated by the PLN/SERCA2 pathway in the TRN are currently under investigation by our lab. Of note, our findings may have clinical neurobiological implications for carriers of human PLN mutations, including the L39stop PLN null mutation that leads to lethal dilated cardiomyopathy [18, 21, 72]. As disruption of the TRN-thalamocortical circuits have been implicated in the pathophysiology of debilitating brain diseases, including schizophrenia, ADHD and ASD [60, 62, 73], gaining insights into the regulation of Ca^2+^ homeostasis in the TRN may hold great promise for the discovery of innovative pharmacotherapeutic targets for the prevention and treatment of these devastating neurodevelopmental disorders.

## 6. DECLARATIONS

### Availability of data and materials

All data supporting the findings of this study are available within the article and supplementary materials, and from the corresponding author on reasonable request.

### Competing interests

The authors declare that they have no competing interests.

### Funding

BK, AB and CTh were supported by the University of Dayton (UD) Graduate School and by the UD Office for Graduate Affairs through the Graduate Student Summer Fellowship (GSSF) Program. KK was supported by the UD Graduate School and by the Department of Biology. JS was supported by a Barry Goldwater Scholarship Award, by the UD Honors Program, by a Lancaster- McDougall award from the Department of Biology, a College of Arts & Sciences (CAS) Dean’s Summer Research fellowship and by UD’s STEM Catalyst Grant program. KB was supported by CAS Dean’s Summer Research fellowships. HO was supported by the UD Honors Program, a CAS Dean’s Summer Research fellowship, and a Lancaster-McDougal departmental fellowship. DS and EGK were supported by a grant from the Leducq Foundation for Cardiovascular Research (CURE-PLaN, 18CVD01). This work was supported by funding from the National Institute of Neurological Disorders and Stroke (NINDS) of the National Institutes of Health (NIH) under award number R03NS109836, as well as by a UD inaugural STEM Catalyst grant, UD Research Council SEED Grants and start-up funding from UD to PMP. Funding sponsors had no further role in study design, in the collection, analysis and interpretation of data, in the writing of the report, and in the decision to submit the article for publication. The construction of the floxed PLN mouse line was funded by an Inaugural STEM Catalyst grant from the University of Dayton to Dr. Pitychoutis.

### Authors’ contributions

BK and AB contributed equally. Drafting of the first manuscript: BK, PMP. Acquisition, analysis and/or interpretation of data: BK, AB, JS, HO, KK, KB, CTh, CTz, DS, EGK, PMP. Study design, supervision and funding acquisition: PMP. All authors reviewed and approved the final version of the manuscript.

## Acknowledgements

The authors would like to thank the undergraduate student members of the Pitychoutis lab, and specifically Emily Flaherty, Kaitlyn Martin, Augustine Miller and Jason Tornes, for assisting with preliminary behavioral experiments and with mouse colony maintenance, and Dr. Dimitrios Arvanitis (Biomedical Research Foundation of the Academy of Athens, Athens, Greece) for his insights into the mRNA assessments. We would like to thank Dr. Hu and the members of the Transgenic Animal and Genome Editing Core at Cincinnati Children’s Hospital Medical Center (CCHMC, Cincinnati, OH) for their service in generating the floxed PLN mouse. The authors would also like to thank Dr. James P. Herman (University of Cincinnati, Cincinnati, OH) for his valuable advice and support with the construction of the conditional mouse line, as well as Dr. Teresa Reyes (University of Cincinnati, Cincinnati, OH) and the staff at Lafayette Instrument (IN, USA) for their support with the 5-SCRTT task.

## Notes

### Competing Interest Statement

The authors have declared no competing interest.

