## Supplementary Table S1 for "A Novel Role for Phospholamban in the Thalamic Reticular Nucleus"

**Supplementary Table S1: Statistically significant results of statistical analyses for behavioral experiments.**

| Paradigm or assay | Parameter measured | N/ genotype/sex | Statistical test | Comparison | Statistics | Deg. freed. | P value | Fig. |
|---|---|---|---|---|---|---|---|---|
| **OFT** | Distance travelled | 17-24 | Three-way RM ANOVA | Main effect of genotype | F=11.201 | 1, 75 | <0.001 | **4a** |
| | | | | Main effect of time | F=120.229 | 4.999, 347.947 | <0.001 | |
| | Time in center | 17-24 | Kruskal-Wallis | *Pln*$^{+/+}$ vs. *Pln*$^{-/-}$ | | | 0.047 | **4b** |
| **Y-maze** | % Spontaneous Alternation score | 17-24 | Two-way ANOVA | Main effect of genotype | F=7.887 | 1, 30 | 0.009 | **4c** |
| **OLM** | Interaction time with objects | 6-12 | 2-way ANOVA | Main effect of object | F=27.19 | 1,72 | <0.001 | **4d** |
| **ORM** | Interaction time with objects | 6-12 | 2-way ANOVA | Main effect of object | F=11.14 | 1,72 | 0.001 | **4e** |
| | | | | Object x genotype interaction | F=4.411 | 1,72 | 0.039 | |
| | | | Bonferroni's multiple comparisons | *Pln*$^{+/+}$: Familiar vs Novel | | | 0.0029 | |
| **3-chamber social interaction test: Social I vs empty** | Interaction time | 5 | Two-way ANOVA | Main effect of stimulus | F=24.18 | 1, 36 | <0.001 | **4f** |
| **3-chamber social interaction test: Social I vs Social II** | Interaction time | 5 | Two-way ANOVA | Main effect of stimulus | F=19.37 | 1, 36 | <0.001 | **4g** |
| **OFT** | Distance travelled (5min bins) | 11-16 | Three-way RM ANOVA | Main effect of time | F=40.085 | 3.717, 193.281 | <0.001 | **5b** |
| | | | | Time x genotype interaction | F=6.685 | 3.717, 193.281 | <0.001 | |
| | Distance travelled (15min) | 11-16 | One-way ANOVA | Main effect of genotype | F=7.403 | 1, 52 | 0.009 | **5c** |
| **EEG Sleep Analysis** | Total: REM duration | 3-5 | Two-way ANOVA | Sex x genotype interaction | F=9.248 | 1, 16 | 0.01 | **6c** |
| | | | Bonferroni's multiple comparisons | Control ♀ vs cKO ♀ | | | 0.048 | |

| | | | | | | | | |
|---|---|---|---|---|---|---|---|---|
| | Dark period: Wakefulness duration | 3-5 | Two-way ANOVA | Main effect of genotype | F=6.171 | 1,16 | 0.029 | **6d** |
| | Dark period: Wakefulness bout length | 3-5 | Two-way ANOVA | Main effect of genotype | F = 8.068 | 1, 12 | 0.0149 | **6d** |
| | Dark period: NREM duration | 3-5 | Two-way ANOVA | Main effect of genotype | F=5.291 | 1, 16 | 0.04 | **6e** |
| | Dark Period:No. of NREM bouts | 3-5 | Two-way ANOVA | Main effect of genotype | F = 4.805 | 1, 12 | 0.0488 | **6e** |
| | Dark period: REM duration | 3-5 | Two-way ANOVA | Main effect of genotype | F=3.481 | 1, 16 | 0.024 | **6f** |
| | | | | Sex x genotype interaction | F=17.934 | 1, 16 | 0.001 | |
| | | | Bonferroni's multiple comparisons | Control ♀ vs cKO ♀ | | | 0.002 | |
| | Dark Period: No of REM bouts | 3-5 | Two-way ANOVA | Sex x genotype interaction | F = 9.937 | 1,12 | 0.0083 | **6f** |
| | | | Bonferroni's multiple comparisons | Control ♀ vs cKO ♀ | | | 0.0494 | |
| | | | | control ♀ vs control ♂ | | | 0.0294 | |
| | Dark Period: REM bout length | 3-5 | Two-way ANOVA | Main effect of genotype | F = 8.458 | 1, 12 | 0.0131 | **6f** |
| | | | | Sex x genotype interaction | F = 8.887 | 1, 12 | 0.0115 | |
| | | | Bonferroni's multiple comparisons | Control ♀ vs cKO ♀ | | | 0.0089 | |
| | Dark Period: Wake-NREM transitions | 3-5 | Two-way ANOVA | Main effect of genotype | F=5.430 | 1, 12 | 0.0381 | **6g** |
| | Dark Period: NREM-Wake transitions | 3-5 | Two-way ANOVA | Main effect of genotype | F = 4.206 | 1, 12 | 0.0628 | **6h** |
| | Dark Period: No. of NREM arousals | 3-5 | Two-way ANOVA | Main effect of genotype | F = 4.677 | 1, 12 | 0.0515 | **6i** |

| | | | | | | | | |
|---|---|---|---|---|---|---|---|---|
| **5-CSRTT: FR1** | FR1 acquisition | 5-9 | Mantel-Cox Log-rank test | cKO ♀ vs ♂ cKO | $x^2$=5.753 | 3 | 0.016 | **7b** |
| | | | | cKO ♀ vs control ♂ | $x^2$=5.560 | 3 | 0.018 | |
| | | | | cKO ♀ vs control ♀ | $x^2$=3.171 | 3 | 0.075 (trend) | |
| **5CSRTT: Titration** | Premature response rate | 5-9 | Two-way ANOVA | Main effect of genotype | F=4.052 | 1, 24 | 0.055 (trend) | **7j** |
| | Premature responses (total) | 5-9 | Two-way ANOVA | Main effect of genotype | F=4.273 | 1, 24 | 0.05 | **7k** |
