## Supplementary Table S2 for "A Novel Role for Phospholamban in the Thalamic Reticular Nucleus"

**Supplementary Table S2: Conditional deletion of *Pln* in the TRN does not affect spectral EEG profile:** No differences detected in EEG spectral profile in delta, theta, beta, alpha, or gamma frequency bands during wakefulness, NREM, or REM sleep when assessing the total, light, or dark period. Data is expressed as power ($\mu V^2$) of delta (δ; 0.5-4Hz), theta (θ; 4-8Hz), alpha (α; 8-13Hz), beta (β; 13-30Hz), gamma (γ; 30-40Hz), frequency bands, normalized as a percentage of total power across all frequency bands.

| | Total | | Light Period | | Dark Period | |
|---|---|---|---|---|---|---|
| | **Control** | **cKO** | **Control** | **cKO** | **Control** | **cKO** |
| *Wakefulness (% of total power)* | | | | | | |
| **Delta** | 52.389±5.414 | 45.146±2.692 | 55.906±5.587 | 50.207±2.327 | 50.412±5.286 | 42.171±2.986 |
| **Theta** | 30.923±3.454 | 35.568±2.06 | 29.189±3.318 | 33.236±2.039 | 31.876±3.478 | 36.922±2.11 |
| **Alpha** | 9.513±1.389 | 10.487±0.844 | 7.912±1.432 | 8.664±0.607 | 10.335±1.396 | 11.557±1.019 |
| **Beta** | 7.148±1.575 | 8.751±1.236 | 6.978±1.737 | 7.84±0.84 | 7.341±1.519 | 9.306±1.493 |
| **Gamma** | 0.027±0.015 | 0.048±0.021 | 0.015±0.007 | 0.053±0.019 | 0.035±0.022 | 0.044±0.023 |
| *NREM Sleep (% of total power)* | | | | | | |
| **Delta** | 55.219±5.864 | 44.586±2.516 | 55.432±6.22 | 44.611±2.759 | 53.239±4.305 | 44.694±2.348 |
| **Theta** | 25.059±2.519 | 29.456±1.117 | 24.322±2.54 | 28.904±1.238 | 27.169±1.884 | 30.498±1.047 |
| **Alpha** | 10.168±1.933 | 12.901±0.976 | 10.443±2.062 | 13.486±1.019 | 10.055±1.587 | 11.931±0.898 |
| **Beta** | 9.55±1.89 | 13.054±1.788 | 9.798±2.013 | 12.994±1.605 | 9.535±1.531 | 12.873±2.125 |
| **Gamma** | 0.004±0.001 | 0.004±0.001 | 0.004±0.001 | 0.004±0.002 | 0.003±0.001 | 0.004±0.001 |
| *REM Sleep (% of total power)* | | | | | | |
| **Delta** | 38.709±5.149 | 29.142±2.149 | 39.075±5.452 | 28.625±2.009 | 36.628±3.69 | 30.155±2.374 |
| **Theta** | 37.571±2.854 | 42.951±1.957 | 36.952±2.942 | 42.956±1.84 | 39.799±2.317 | 42.82±2.401 |
| **Alpha** | 12.686±1.72 | 14.129±0.755 | 12.707±1.765 | 14.443±0.639 | 12.977±1.556 | 13.611±0.951 |
| **Beta** | 11.029±1.495 | 13.772±2.301 | 11.26±1.515 | 13.97±2.135 | 10.593±1.433 | 13.409±2.754 |
| **Gamma** | 0.004±0.001 | 0.005±0.002 | 0.005±0.001 | 0.005±0.002 | 0.003±0.001 | 0.005±0.002 |
