## Supplementary figures and images for "A Novel Role for Phospholamban in the Thalamic Reticular Nucleus"

### Supplementary Figure S1

#### Supplementary Figure S1 (Fig. S1)

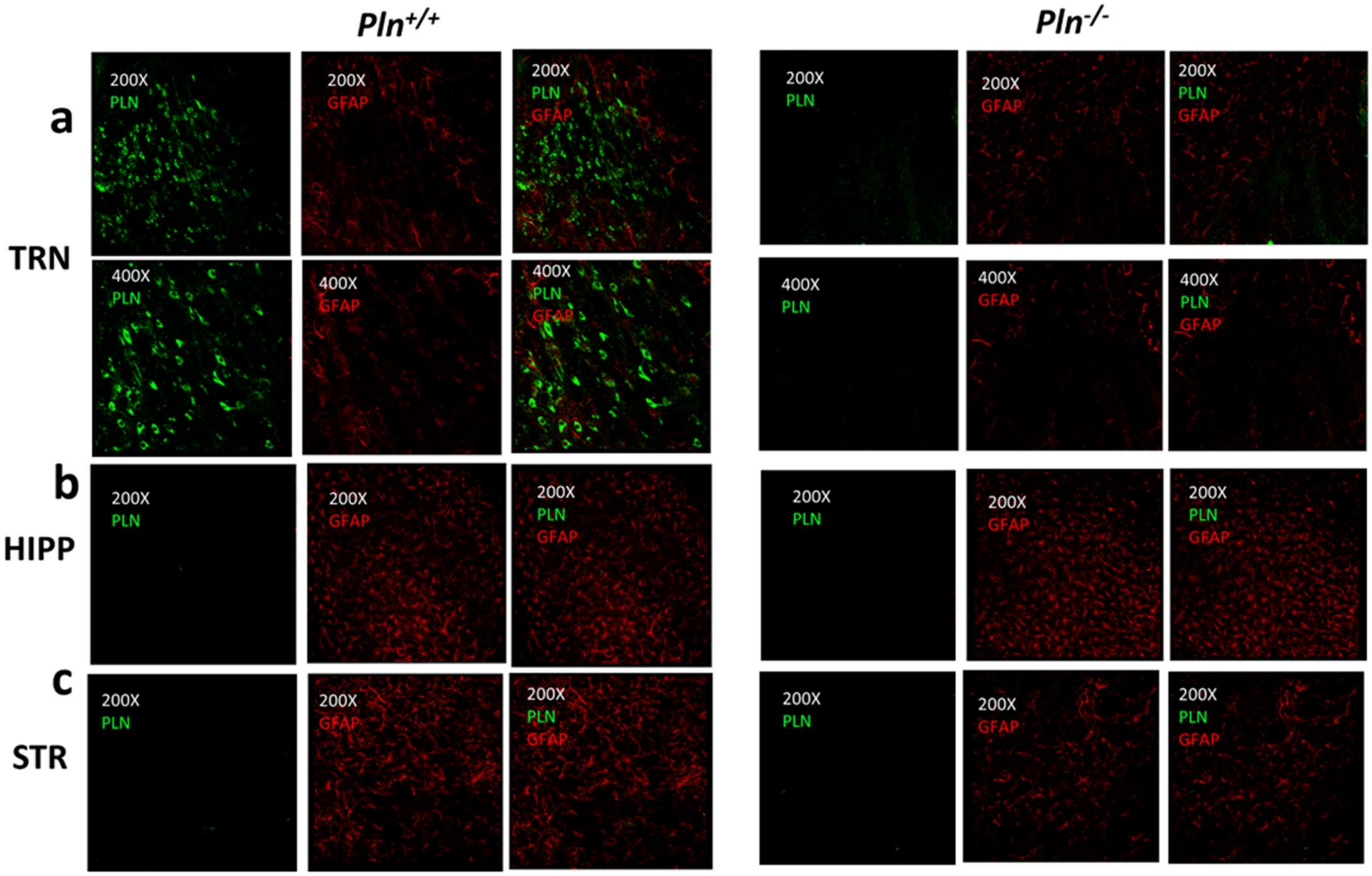
